# SRY: An Effective Method for Sorting Long Reads of Sex-limited Chromosome

**DOI:** 10.1101/2020.05.25.115592

**Authors:** Xiao-Bo Wang, Qing-You Liu, A-Lun Li, Jue Ruan

## Abstract

Most of available reference genomes are lack of the sequence map of sex-limited chromosomes, that make the assemblies uncompleted. Recent advances on long reads sequencing and population sequencing raise the opportunity to assemble sex-limited chromosomes without the traditional complicated experimental efforts. We introduce a computational method that shows high efficiency on sorting and assembling long reads sequenced from sex-limited chromosomes. It will lead to the complete reference genomes and facilitate downstream research of sex-limited chromosomes.

## Main

Generally, most of genome projects prefer homogametic (XX females or ZZ males) to heterogametic (XY males or ZW females) genomes sequencing, because latter haploid nature of both sex chromosomes (X and Y, or Z and W) provide reduced sequencing depth that can decrease assembled contiguity and length^1^. Besides, as XY or ZW chromosomes were evolved from a pair of autosomal chromosomes^2, 3^, such homology gives rise to fragmented contigs in the same way with large repeats. Plenty of repetitive sequences in sex-limited (Y or W) chromosome further increase the assembling difficulty and cause the paucity of sex-limited chromosome.

So far, there are principally two approaches aimed to solve the problem. The first, BAC-based method, was once successfully applied on deciphering the Y chromosomes of several mammals including human^3^, chimpanzee^4^, rhesus macaque^5^, and mouse^6^. It employs single-haplotype iterative mapping and sequencing (SHIMS), provides the best solution so far to overcome the difficulty of assembling Y chromosome. Unfortunately, it is time-consuming, labor-intensive and expensive. Afterwards, Chromosome flow-sorting based on chromosome size and GC content was applied. The latter method has high automation and throughput^1^. However, it can mistakenly sort other chromosomes or debris having similar sizes or GC contents with sex-specific chromosome, and bring in biases during the amplification stage^1, 7^.

Thanks to the increasing sequencing read length, long reads have the potential to be identified to its original chromosome by pure computing. Recently, trio binning was developed to utilize specific markers for long reads sorting *in silicon*^8^. It compares length k subsequences (*k*-mers) of short reads from parental genomes and identifies *k*-mers that are unique to each parent. Trio binning further uses these *k*-mers to separate long reads of the diploid offspring and conducts *de novo* haplotype assemblies, separately. Theoretically, Y- (or W-) specific markers can be selected and used for sorting long reads from sex-limited chromosome. Compared to whole genome assembly (WGA), the trio binning assembly covers ~2.1Mb more genomic regions of Y chromosome with a comparable contiguity (Table 1). It indicates that computational method is promising, though trio binning cannot efficiently address the problem of assembling sex-limited chromosome based on its scheme to select specific markers.

**Table 1.** Comparison of assembled genomes from SRY, trio binning and WGA.

| Method | SRY | SRY | SRY | SRY | Trio binning | WGA |
| --- | --- | --- | --- | --- | --- | --- |
| Assembly software | Shasta | Canu | Flye | Wtdbg2 | Wtdbg2 | Wtdbg2 |
| GRch38 Y length | 57.2Mb | 57.2Mb | 57.2Mb | 57.2Mb | 57.2Mb | 57.2Mb |
| Assembled contig length | 19.5Mb | 24.5M | 20.8Mb | 23.7Mb | 14.4Mb | 12.0Mb |
| Largest contig | 4,512,408 | 6,054,034 | 5,141,512 | 6,372,558 | 2,107,873 | 2,082,680 |
| N50 | 2,617,545 | 3,275,858 | 2,483,958 | 1,494,959 | 1,386,955 | 1,440,510 |
| Aligned length | 18.9Mb | 22.4Mb | 19.5Mb | 19.4Mb | 13.8Mb | 11.7Mb |
| NA50 (50% query in<br>alignments longer than<br>NA50) | 1,084,084 | 1,019,072 | 1,162,224 | 1,076,192 | 1,068,368 | 1,076,825 |
| NA75 (75% query in<br>alignments longer than<br>NA75) | 328,410 | 231,009 | 309,942 | 44,041 | 239,262 | 239,233 |
| No. of mismatches per 100<br>kbp | 90.63 | 399.42 | 171.03 | 157.54 | 129.04 | 149.59 |
| No. of indels per 100 kbp | 254.23 | 701.98 | 270.97 | 647.34 | 639.60 | 659.02 |

The nature of difference between sex chromosomes and autosomes in whole genome sequencing is half of the read depth. In heterogametic (such as XY male) populations, sex chromosomes consist of X and Y chromosome, and the sequencing depth of each chromosome to autosome is half. In homogametic (such as XX female) populations, sex chromosome only consists of X chromosome. X/Y-specific markers of male populations can be separated depending on depth. Because only X-specific markers exists in female populations, they can be excluded from X/Y-specific markers to identify Y-specific markers.

To reach the goal of sorting long reads of sex-limited chromosome, we developed an *in silicon* sorting method called SRY. SRY firstly selects half-depth *k*-mers which in plot located in first peak (hybrid peak) of the *k*-mer depth distribution of target male individual sequencing (Fig. 1a). Then, it filters X-linked *k-*mers included in female populations (Fig. 1b). Besides, SRY filters *k*-mers of autosomes that originate from heterozygosity to identify male-specific *k-*mers (MSK) (Fig. 1b). Owing to the impact of population structure and sequencing errors, the operation of SRY is in fact a sampling process, which unavoidably involves discrete *k*-mers from X chromosome and autosomes. So SRY calculates MSK density of long reads and excludes those with lower marker density (Fig. 1c).

**Figure 1.**
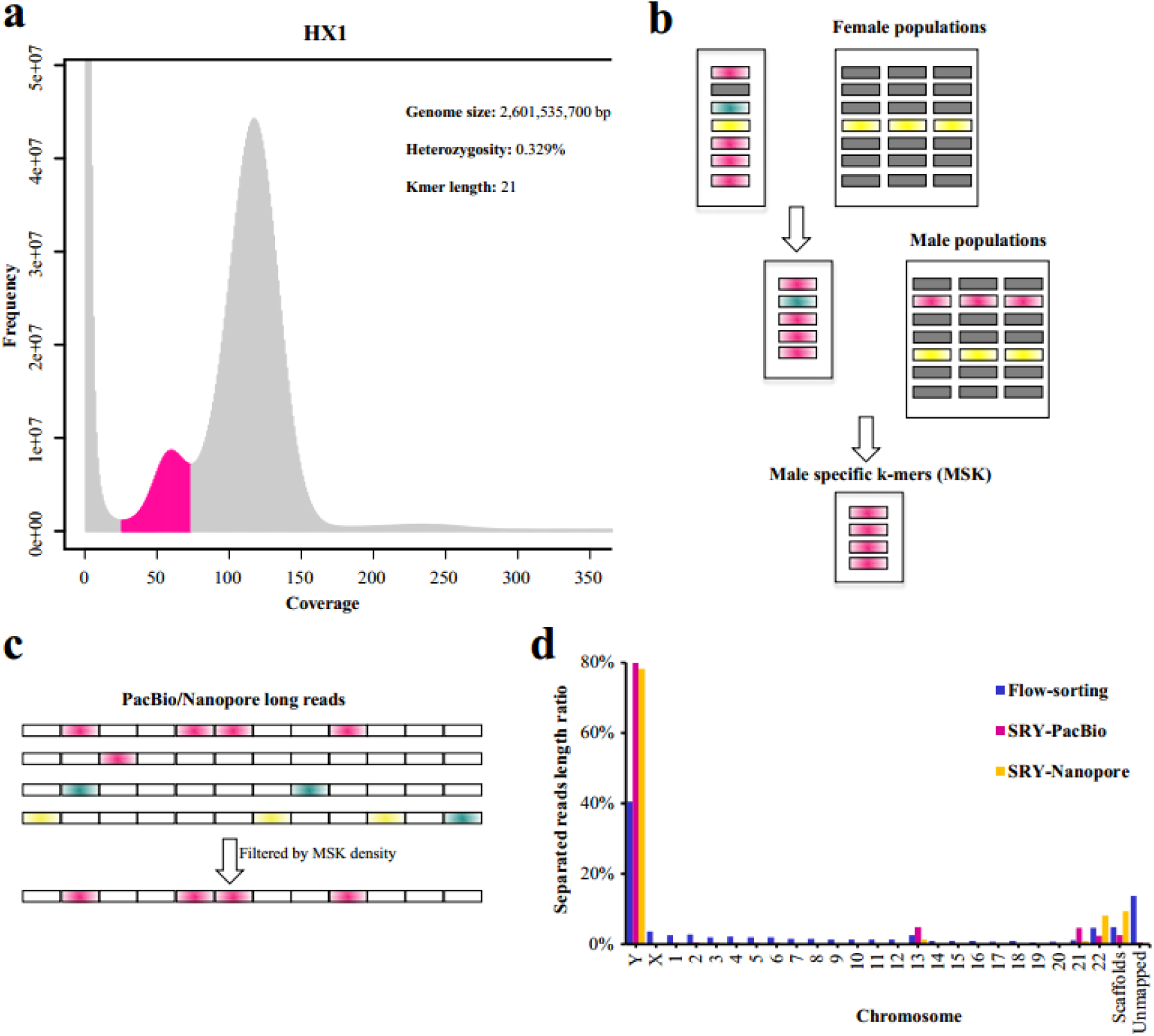
The strategy of SRY for sorting long reads. (**a**) The distribution of *k-*mers of HX1 short reads was plotted using GenomeScope. *K-*mers in the first peak (in red) were selected for the initial collection of HX1 *k-*mers. (**b**) SRY firstly removed those contained in female populations from HX1 *k-*mers. Then, SRY retained *k-*mers that occurred in least one individual of male populations. Red rectangle represents male specific *k-*mer (MSK), and yellow rectangle represents X-linked *k-*mers, and green rectangle represents *k-*mers of HX1 autosomes that originate from heterozygosity. Those colored by gray represent common *k-*mers in both male and female populations. (**c**) SRY selected PacBio/Nanopore long reads whose MSK density higher than the average value. The meanings of colors are in accordance with **b**. (**d**) Long reads sorted by SRY and flow-sorting were mapped on GRch38 reference genome, separately. SRY shows significantly higher sorting efficiency on Y chromosome.

To evaluate the sorting efficiency of SRY, we downloaded datasets including short and long reads of a Chinese individual HX1^9, 10^, and whole genome sequencing short reads of a Han Chinese population (44 females and 46 males)^11^ (**Supplementary Table 1 and Table 2**). SRY obtained about 10 million MSK as well as sorted ~3.7Gb (71X) PacBio and Nanopore long reads of Y chromosome (**Supplementary Table 3**). We used minimap2^12^ to align these sorted long reads and Nanopare data from flow-sorting method^7^ on GRch38 reference genome, separately. Whether on sorting Nanopare reads or PacBio reads, SRY showed nearly one times higher efficiency than the flow-sorting method (Fig. 1d).

**Table 2.**
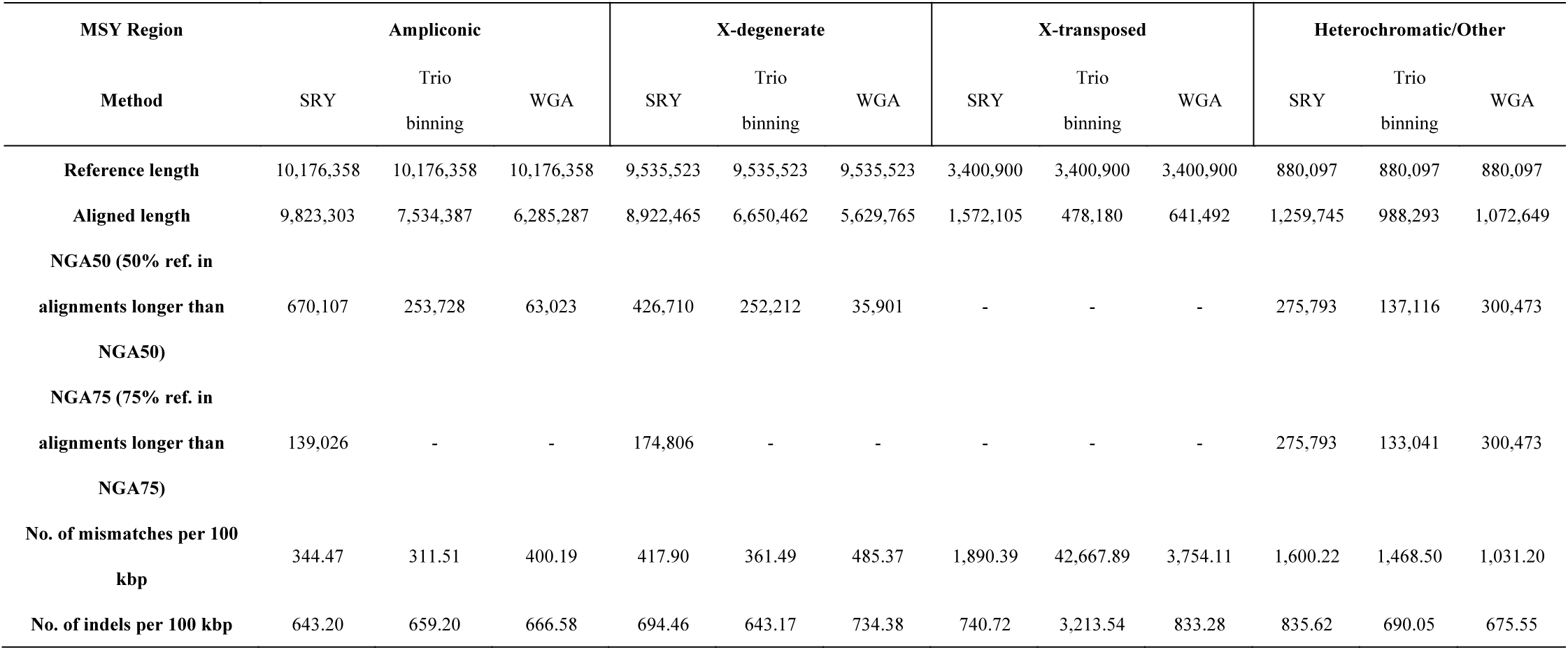
Performance of three methods on four discrete regions of the GRch38 MSY. “-” indicates that the metric is incalculable.

To assess the assembling completeness of Y chromosome with and without specific markers, we compared the resulting assembly from SRY, trio binning and WGA using a father-mother-offspring trio dataset^13^. SRY used 10 million MSK above and sorted ~1.3Gb (25X) Nanopore reads of the offspring (**Supplementary Table 3**). We assembled those sorted reads with popular software including wtdbg2^14^, flye^15^, canu^16^ and shasta^13^. All these assemblers provide contig N50 of at least 1.5Mb (Table 1). Canu that supports for trio binning was used to separate long reads of male-haplotype. We applied the fast assembler wtdbg2 to assemble those sorted reads and conduct whole genome assembly. The total size of genome from SRY is about 2 times as long as those from both trio binning and WGA (Table 1). Its contiguity and assembly accuracy are comparable to other two methods. SRY provides longest aligned length on all four class of MSY (male-specific region of Y chromosome), and largest NGA50 in ampliconic and X-degenerate regions (Table 2). In the X-transposed region, which exhibits 99% identity to the X chromosome, the aligned length from SRY is 2-4 times longer than others. Furthermore, SRY shows 2 to 30 times lower density of mismatches and indels in this region (Table 2). The heterochromatic sequences fulfill abundant tandem repeats of low sequence complexity, and thus may cause misalignments and aligned lengths of all three methods longer than the reference.

Complete *de novo* assembly of sex-limited chromosome based on whole-genome sequencing is challenging due to its haploid nature, homology to X/Z chromosome and highly repetitive contents, hindering efforts to construct full-length genome and understand sex-specific morphology and evolution. SRY sorts long reads of sex-limited chromosome and can improve the completeness of assembling result. Nowadays, a genome sequencing project always contains a deep individual sequencing for reference genome and re-sequencing for populations. The advent of SRY offers the opportunity to choose a male individual and construct the reference sequence for all chromosomes. With highly accurate long reads being available^17^, SRY will be more promising to assemble the completed reference genomes.

## Methods

### SRY process

Firstly, SRY uses kmer_count to acquire 21-mer sets from short reads of targeted male species and populations. Next, the program filterx is used to remove *k-*mers of targeted species that appear in female populations. The collection of *k-*mers are further filtered by selecting those included in at least one individual of male populations. Then, SRY selects long reads of targeted species that have male specific *k-*mers (MSK). Finally, SRY filters those long reads with lower MSK densities than average value of whole Y chromosome. The average MSK density per 1kb is calculated as follow:

*T*(1-E)21*1000/G*

T represents the total number of MSK, and E represents the base error rate of long reads (15%) and G indicates genome size of Y chromosome (52M for human Y).

SRY identified MSK of HX1 with the command “SRY -s HX1.fq.gz -m male-population.fq.gz -f female-population.fq.gz -l HX1.long-reads.fasta.gz -g 52 -i 27”.

### Assessment

We simulated 50X Nanopore and PacBio reads (25X for Y chromosome) based on GRch38 genomes using badread^18^ package (v0.1.3) with the following commands, respectively: *badread simulate --reference GRch38_autoX.fa --quantity 50X --error_model nanopore -- start_adapter 0,0 --end_adapter 0,0 --junk_reads 0 --random_reads 0 --chimeras 0* (simulated Nanopore reads of autosomes and X chromosome)

*badread simulate --reference GRch38_Y.fa --quantity 25X --error_model nanopore --start_adapter 0,0 --end_adapter 0,0 --junk_reads 0 --random_reads 0 --chimeras 0* (simulated Nanopore reads of Y chromosome)

*badread simulate --reference GRch38_autoX.fa --quantity 50X --error_model pacbio --identity 85,95,3 --length 7500,7500 --start_adapter 0,0 --end_adapter 0,0 --junk_reads 0 --random_reads 0 --chimeras 0* (simulated PacBio reads of autosomes and X chromosome)

*badread simulate --reference GRch38_Y.fa --quantity 25X --error_model pacbio --identity 85,95,3 --length 7500,7500 --start_adapter 0,0 --end_adapter 0,0 --junk_reads 0 --random_reads 0 -- chimeras 0* (simulated PacBio reads of Y chromosome)

A total sum/number of 142.7Gb/10,024,130 Nanopore reads (557.3Mb/39,452 for Y chromosome) and 113.8Gb/15,564,739 PacBio reads (574.7Mb/78,830 for Y chromosome) were generated, excluding those contained N in sequences. Depending on these datasets, we evaluated the specificity and sensitivity of SRY using MSK identified from HX1. The filtering threshold with average MSK density (6.3/kb) had a good performance on sorting Nanopore reads, reaching about 100% specificity and 83.3% sensitivity. As expected, the sensitivity for PacBio reads is relatively decreased, reaching 66.9%.

### Y chromosome assembly and identification

We collected the dataset of a trio family including short reads from father (HG01107, ~113X) and mother (HG01108, ~79X) and Nanopore reads from child (HG01109, ~72X)^13^. SRY separated 1.3G (~25X) long reads of HG01109 using MSK markers identified from HX1. Then, we used wtdbg2.5 with parameters “-L 0 -p 0 -k 21 -s 0.25 -S 2 --rescue-low-cov-edges” to assembly those long reads. We also used other assemblers including Canu (v2.0), Flye (v2.6) and Shasta (v0.4.0) with default parameters to perform the genome assembly. Trio binning phased HG01109 long reads using with command “canu - stopAfter=haplotype genomeSize=3g -haplotypeMale HG01107.fastq.gz -haplotypeFemale HG01108.fastq.gz -nanopore-raw HG01109.fasta.tar.bz2”. To compare with SRY, wtdbg2 with same parameters (except for the genome size parameter “-g 3.1G”) was applied to assembly phasing reads from trio binning and conduct whole genome assembly (WGA).

To identify Y genome of trio binning and WGA, we used nucmer of MUMmer^19^ package (version 4.0.0beta2) with default parameter to align their assemblies with GRch38 reference. Delta-filter with parameter “-i 95” was further used to filter the alignment results. Finally, the best-hit alignments with at least 90% coverage on reference Y chromosome was retained. We utilized quast (v5.0.2) with default parameters to evaluate the assembled genome quality.

## Data availability

We downloaded all Nanopore, PacBio and Illumina datasets under NCBI project number PRJNA301527 for HX1. The SRA numbers of Han Chinese population were listed at supplementary table 1 and 2. The trio family (HG01107, HG01108 and HG01109) reads are available at https://s3-us-west-2.amazonaws.com/human-pangenomics/index.html. We also downloaded ERR3241824 for HG01107 and ERR3241825 for HG01108 to improve the performance of trio binning.

## Code availability

The SRY source code is hosted by GitHub at: https://github.com/caaswxb/SRY.

## Acknowledgments

This work was supported by National Natural Science Foundation of China (Grant No. 91731304 to J.R.), National Key Research and Development Program of China (Grant No. 2019YFA0707003 to J.R.) and National Natural Science Foundation of China (Grant No. 31860638 to Q. L.). We thank S. Wu from CAAS for his suggestions on genome assembly.

## Author contributions

J.R. and Q. L. designed the project, and J.R. managed the project. X.W., J.R. and A.L. developed the SRY method. X.W. collected genomic data, performed analysis and wrote the paper. J.R. revised the manuscript.

## Competing interests

The authors declare no competing interests.

## Notes

### Competing Interest Statement

The authors have declared no competing interest.

## Reference

1. Tomaszkiewicz, M., Medvedev, P. & Makova, K.D. Y and W Chromosome Assemblies: Approaches and Discoveries. Trends in Genetics 33, 266–282 (2017).

2. Bellott, D.W. et al. Avian W and mammalian Y chromosomes convergently retained dosage-sensitive regulators. Nat. Genet. 49, 387–394 (2017).

3. Skaletsky, H. et al. The male-specific region of the human Y chromosome is a mosaic of discrete sequence classes. Nature 423, 825–837 (2003).

4. Hughes, J.F. et al. Chimpanzee and human Y chromosomes are remarkably divergent in structure and gene content. Nature 463, 536–539 (2010).

5. Hughes, J.F. et al. Strict evolutionary conservation followed rapid gene loss on human and rhesus Y chromosomes. Nature 483, 82–86 (2012).

6. Soh, Y.Q.S. et al. Sequencing the Mouse Y Chromosome Reveals Convergent Gene Acquisition and Amplification on Both Sex Chromosomes. Cell 159, 800–813 (2014).

7. Kuderna, L.F.K. et al. Selective single molecule sequencing and assembly of a human Y chromosome of African origin. Nat. Commun. 10, 1–8 (2019).

8. Koren, S. et al. De novo assembly of haplotype-resolved genomes with trio binning. Nat. Biotechnol. 36, 1174–1182 (2018).

9. Shi, L. et al. Long-read sequencing and de novo assembly of a Chinese genome. Nat. Commun. 7, 1–10 (2016).

10. Liu, Q. et al. Detection of DNA base modifications by deep recurrent neural network on Oxford Nanopore sequencing data. Nat. Commun. 10, 1–11 (2019).

11. Lan, T. et al. Deep whole-genome sequencing of 90 Han Chinese genomes. Gigascience 6, 1–7 (2017).

12. Li, H. Minimap2: pairwise alignment for nucleotide sequences. Bioinformatics 34, 3094–3100 (2018).

13. Shafin, K. et al. Nanopore sequencing and the Shasta toolkit enable efficient de novo assembly of eleven human genomes. Nat. Biotechnol. (2020).

14. Ruan, J. & Li, H. Fast and accurate long-read assembly with wtdbg2. Nat. Methods 17, 155–158 (2020).

15. Kolmogorov, M., Yuan, J., Lin, Y. & Pevzner, P.A. Assembly of long, error-prone reads using repeat graphs. Nat Biotechnol 37, 540–546 (2019).

16. Koren, S. et al. Canu: scalable and accurate long-read assembly via adaptive k-mer weighting and repeat separation. Genome Res. 27, 722–736 (2017).

17. Wenger, A.M. et al. Accurate circular consensus long-read sequencing improves variant detection and assembly of a human genome. Nat. Biotechnol. 37, 1155–1162 (2019).

18. Wick, R. Badread: simulation of error-prone long reads. The Journal of Open Source Software (2019).

19. Kurtz, S. et al. Versatile and open software for comparing large genomes. Genome Biol. 5, R12 (2004).

